# Chromosome III aneuploidy enhances ethanol tolerance in industrial *Saccharomyces cerevisiae* by increasing *TUP1* expression

**DOI:** 10.1101/2025.04.07.647507

**Authors:** Sonia Albillos-Arenal, Javier Alonso del Real, Eladio Barrio, Amparo Querol

## Abstract

Ethanol stress poses a considerable challenge for *Saccharomyces cerevisiae* during fermentation. Strains carrying an extra copy of chromosome III exhibit enhanced ethanol tolerance. Here, we investigated the underlying mechanisms of this tolerance, focusing on gene dosage effects and differential gene expression under ethanol stress. We compared the gene expression profiles of a strain with three copies of chromosome III and its derivative with two copies, exposed to 6% and 10% ethanol. Our analysis identified *TUP1*, located on chromosome III, as a key regulator of the ethanol stress response. Deleting one copy of *TUP1* in the tolerant strain diminished its ethanol tolerance, suggesting that chromosome III aneuploidy in ethanol-tolerant strains enhances adaptive responses by increasing *TUP1* copy number. Our findings offer insights into the genetic basis of ethanol tolerance, with potential applications for optimizing industrial fermentation processes and understanding the role of aneuploidy in the domestication of industrial yeasts.

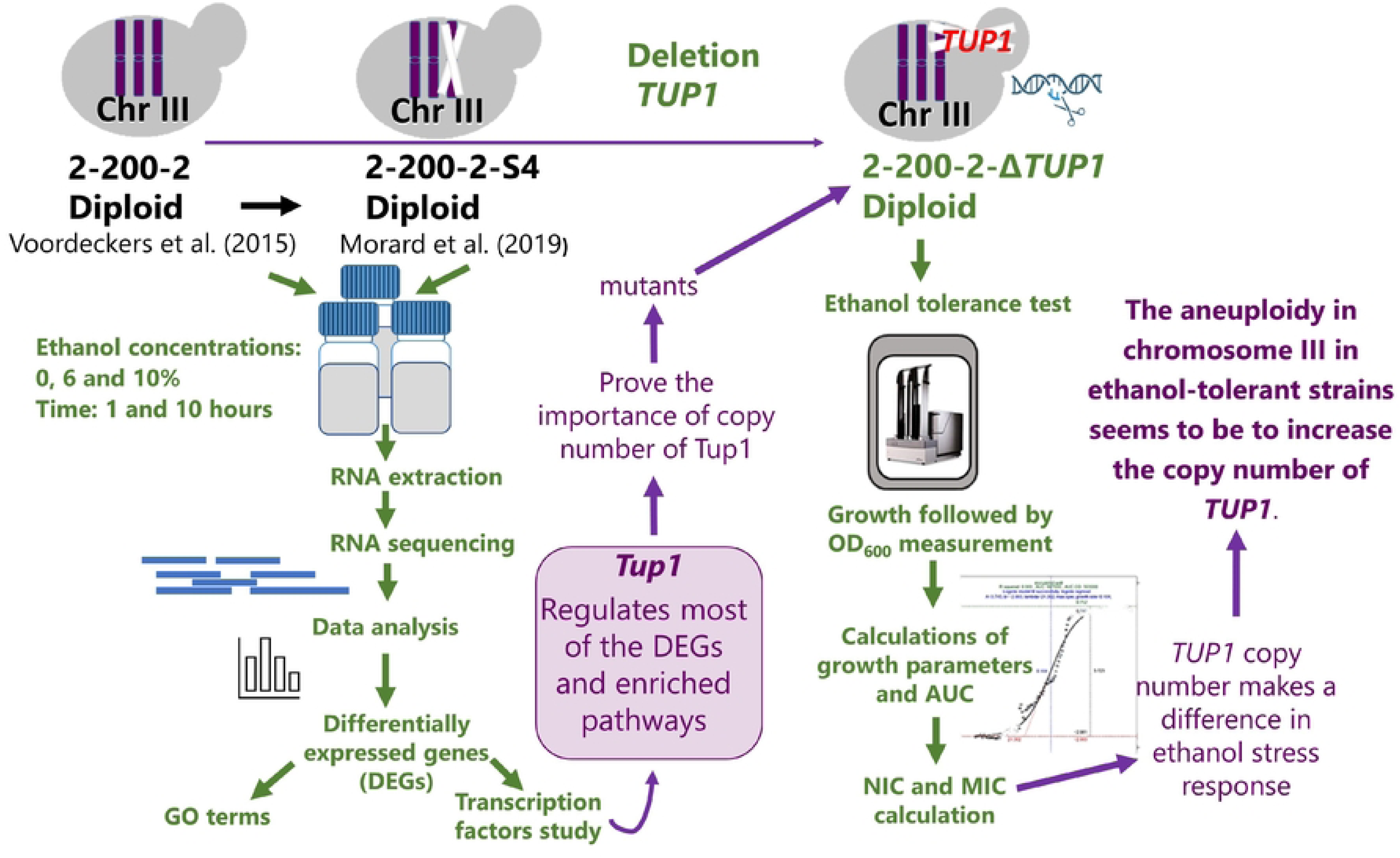

## INTRODUCTION

One key reason for the widespread use of *Saccharomyces* in fermentation is its remarkable tolerance to relatively high ethanol concentrations. However, ethanol remains a potent stressor, impacting numerous cellular processes [1]. Specifically, it inhibits respiration, arrests cell growth, and can ultimately cause cell death [2,3]. To counteract this stress, *S. cerevisiae* activates various defense mechanisms, including the production of heat shock proteins [4], the accumulation of trehalose [5], and changes in vacuole morphology [6]. Additionally, ethanol disrupts protein function, reducing enzyme activity [7,9], compromises membrane integrity [10,11], triggers the production of reactive oxygen species, alters carbon metabolism [12], and affects post-translational modifications such as SUMOylation [13]. These complex responses pose significant challenges to improving fermentation efficiency and sustainability across various industries [14]. Accordingly, developing ethanol-tolerant yeast strains is a key objective in biotechnology. Different strains of *Saccharomyces* exhibit varying levels of tolerance to ethanol stress. Generally, industrial strains such as wine strains [15,16] have a higher tolerance to ethanol stress than wild strains [10]. This is a result of human selection over thousands of years of winemaking, a process known as domestication [17]. *S. cerevisiae* is well-adapted to tolerate ethanol [18], making it a valuable tool in various fermentation processes.

Ethanol tolerance in yeast is a multifaceted process involving numerous genes across the genome. Several mechanisms have been described to mitigate the harmful effects of ethanol [3,8,15,19,20]. Voordeckers et al. (2015) investigated the complex evolutionary adaptations of *S. cerevisiae* to high ethanol concentrations. Through experimental evolution and genomic analysis, they found that aneuploidy of chromosome III significantly contributed to increased ethanol tolerance [21]. This finding was further supported by Morard et al. (2019), who observed a strong correlation between polysomy of chromosome III and high ethanol tolerance [22]. The prevalence of aneuploidy in chromosome III underscores its importance in enhancing ethanol tolerance. Chromosome duplication is a common adaptation strategy for organisms facing harsh environments [23] and acts as an initial defence mechanism, providing a rapid but temporary solution to sudden selective pressures [24].

Interestingly, aneuploidy states can be surprisingly stable in nature, particularly due to protein turnover processes. This contrasts with the transient nature of aneuploidy in laboratory strains [25]. These observations highlight the dynamic interplay between genomic responses to stress and the evolutionary trajectory of organisms [24]. While aneuploidy in chromosome III is linked to increased ethanol tolerance, the underlying mechanisms remain unclear. Understanding this relationship is key to filling knowledge gaps in yeast genetics and potentially improving industrial fermentation processes.

We hypothesize that specific elements on chromosome III are responsible for the beneficial effects observed with increased copy number. One possibility is that key ethanol tolerance genes located on chromosome III are duplicated, leading to an increased gene copy number that results in elevated mRNA transcription and enhanced protein production of these important genes. Another possibility is that a transcription factor situated on chromosome III benefits from the extra copy number, causing differential regulation of its target genes involved in the ethanol stress response. This amplification may ultimately enhance ethanol tolerance. Our analysis revealed differential expression of several genes between two strains with varying copy numbers of chromosome III. However, the most significant finding was the identification of *TUP1*, located on chromosome III, as a crucial regulator of the ethanol stress response. *TUP1* is a global transcriptional repressor that plays a crucial role in regulating various cellular processes in *S. cerevisiae*, including the response to environmental stresses such as ethanol exposure [26,28]. *TUP1* functions in association with *CYC8* (also known as *SSN6*) to form the *TUP1*-*CYC8* complex [26,29,30]. Despite its primary role as a repressor, *TUP1*-*CYC8* complex indirectly activates genes required for the stress response [27]. When we deleted one copy of *TUP1* in the ethanol-tolerant strain, we observed a reduction in its ethanol tolerance. This suggests that the aneuploidy of chromosome III in ethanol-tolerant strains enhances adaptive responses specifically by increasing the copy number of Tup1p. These findings provide valuable insights into the genetic underpinnings of ethanol tolerance, which could have important applications in optimizing industrial fermentation processes. Moreover, our results contribute to the broader understanding of how aneuploidy may have played a role in the domestication of industrial yeast strains.

## RESULTS

### Comparative transcriptome analysis of yeast strains with two or three copies of chromosome III

To investigate the potential link between chromosome III copy number and enhanced ethanol tolerance, we compared the gene expression profiles of two genetically similar *S. cerevisiae* strains under ethanol stress. One strain (2-200-2) carried three copies of chromosome III, while the other had two copies (2-200-2-S4) due to the deletion of one copy [22]. Both strains were cultured in glucose-peptone-yeast extract (GPY) medium containing 0%, 6%, or 10% ethanol, and samples were collected after 1 and 10 hours of exposure for gene expression analysis.

Principal component analysis (PCA) of gene expression levels revealed consistent clustering patterns across the two strains (Fig. S1), suggesting that culture conditions and ethanol exposure time were the determinants of gene expression rather than strain-specific differences. Indeed, inter-strain variations appeared minimal. As the sole genetic distinction between these strains is the copy number of chromosome III, it is plausible that these subtle strain-specific expression differences are primarily driven by this chromosomal variation.

We analyzed RNA-seq data to examine how increased gene dosage on chromosome III affects the regulation of differentially expressed genes (DEGs) and overall transcriptional changes in metabolic pathways. Since few genes showed differential expression without ethanol stress, and had a log fold-change (FC) lower than 2 (Supplementary Table S1), we focused our analysis on DEGs identified after ethanol exposure. Some differentially expressed genes (DEGs) overlapped across different time points and ethanol concentrations (Supplementary Fig. S2). Notably, the highest number of shared genes was observed in the 10% ethanol condition at both 1 and 10 hours of fermentation. To isolate the effects of the chromosomal imbalance, we separately analysed genes on chromosome III and those on other chromosomes (Fig. 1).

**Fig. 1.**
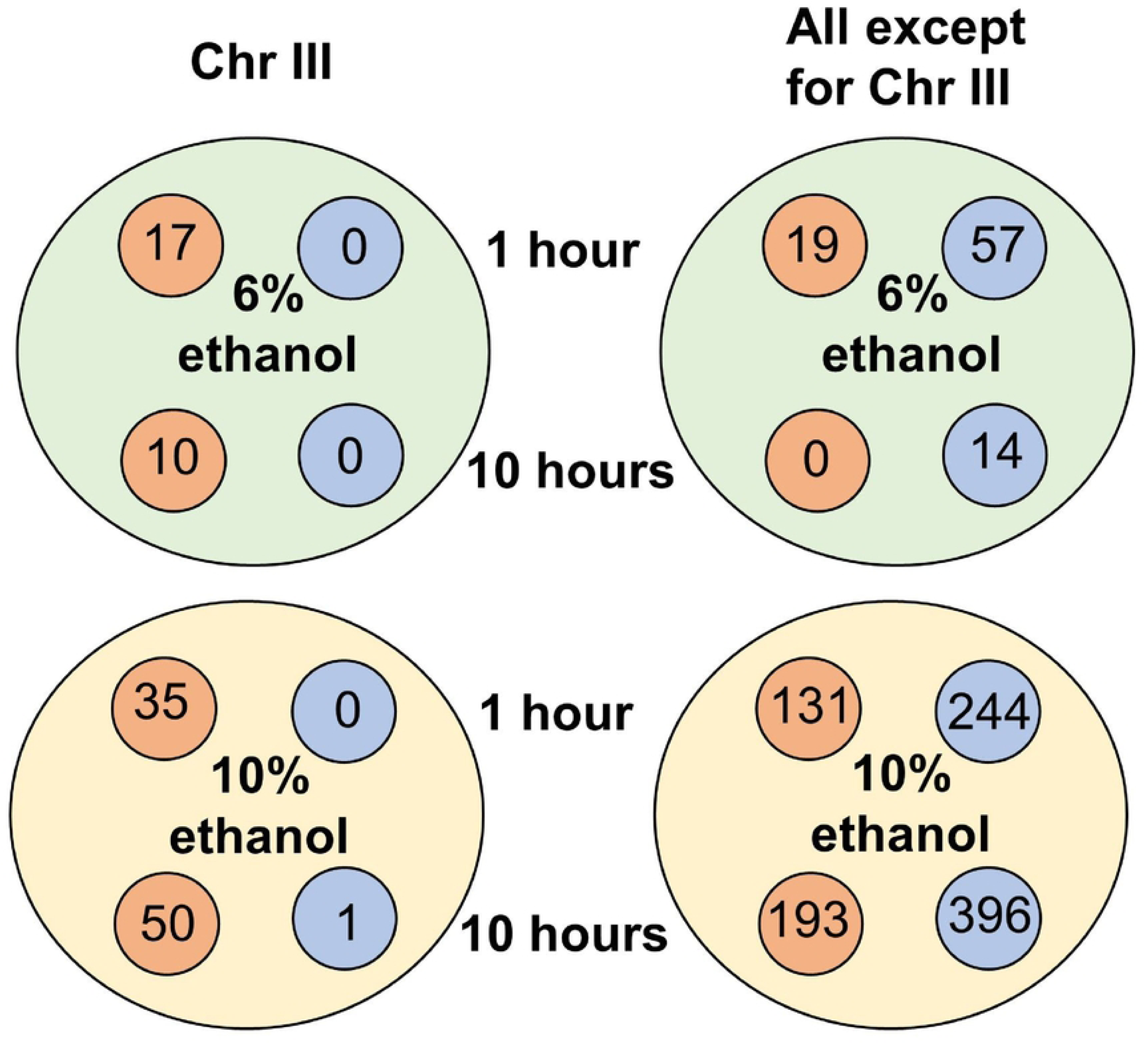
Number of genes up-regulated (orange circles) or down-regulated (blue circles) in yeast exposed to 6% and 10% ethanol, at 1 or 10 hours after the start of fermentation. The figure separately displays the differentially-expressed genes (DEGs) located on chromosome III and those located on other chromosomes.

### Ethanol stress increases the expression of genes located on chromosome III

In this study, our primary objective was to elucidate the mechanisms by which chromosome III aneuploidy contributes to enhanced ethanol tolerance. To achieve this, we focused our analysis on the DEGs located on chromosome III. We hypothesized that genes on this chromosome exhibiting significantly different logFC values between the two strains could play a crucial role in the ethanol response. Expression analysis revealed that 17 genes showed a higher FC (with a p-value cut-off of 0.05) in strain 2-200-2 than in strain 2-200-2-S4 after 1 hour of 6% ethanol exposure, and 10 genes after 10 hours (Fig. 1). When exposed to 10% ethanol, the number of genes increased to 35 after 1 hour and 50 after 10 hours. Notably, 5 genes (*SPB1*, *GFD2*, *ADF1*, *SRO9*, and *RSA4*) exhibited a logFC >2 (Supplementary Table 2). These genes play crucial roles, either directly or indirectly, in ribosome biogenesis or translation regulation, essential processes for protein synthesis. Only one gene, YCR024C-A (*PMPI*), showed lower expression in strain 2-200-2 than in strain 2-200-2-S4 (logFC -1.37). This occurred after 10 hours exposure to 10% ethanol. *PMP1* encodes the regulatory subunit of the plasma membrane H(+)-ATPase Pma1p. This finding suggests that regulatory mechanisms are in place to suppress its expression, even in the presences of an extra copy of the gene.

We then conducted an analysis of DEGs across all chromosomes to identify potential regulators on chromosome III. This investigation aimed to elucidate the factors that could explain the observed differences in gene expression and enriched pathways between strains with varying copies of chromosome III.

### Regulation of differently expressed genes under ethanol stress

As our primary goal was to identify a transcription factor encoded on chromosome III that regulates genes involved in ethanol tolerance, we systematically analyzed the DEGs between the two strains with varying chromosome III copy numbers (Supplementary Table S1) and examined shared regulatory patterns using the DB database [31]. This database contains information on mutant regulator expression, allowing us to investigate transcription factors that regulate these DEGs. Among the identified regulators, the Tup1p-Ssn6p complex emerged as the most prominent candidate with a p-value of < 10^-6^ (Fig. 2). Furthermore, regulator enrichment analysis revealed Tup1p as the sole enriched transcription factor located on chromosome III. Notably, *YCR084C* (*TUP1*) displayed increased expression in the tolerant strain under various ethanol exposure conditions (Supplementary Table S1), with logFC values ranging from 0.64 to 0.81. Additionally, we utilized the YEASTRACT+ database [32], which catalogs regulatory relationships between transcription factors and their target genes, to analyze *TUP1*-regulated genes in each condition (Fig. 3). Significant p-values were observed at 10% ethanol, with the lowest values after 10 hours for overexpressing genes and after 1 hour for repressing genes.

**Fig. 2.**
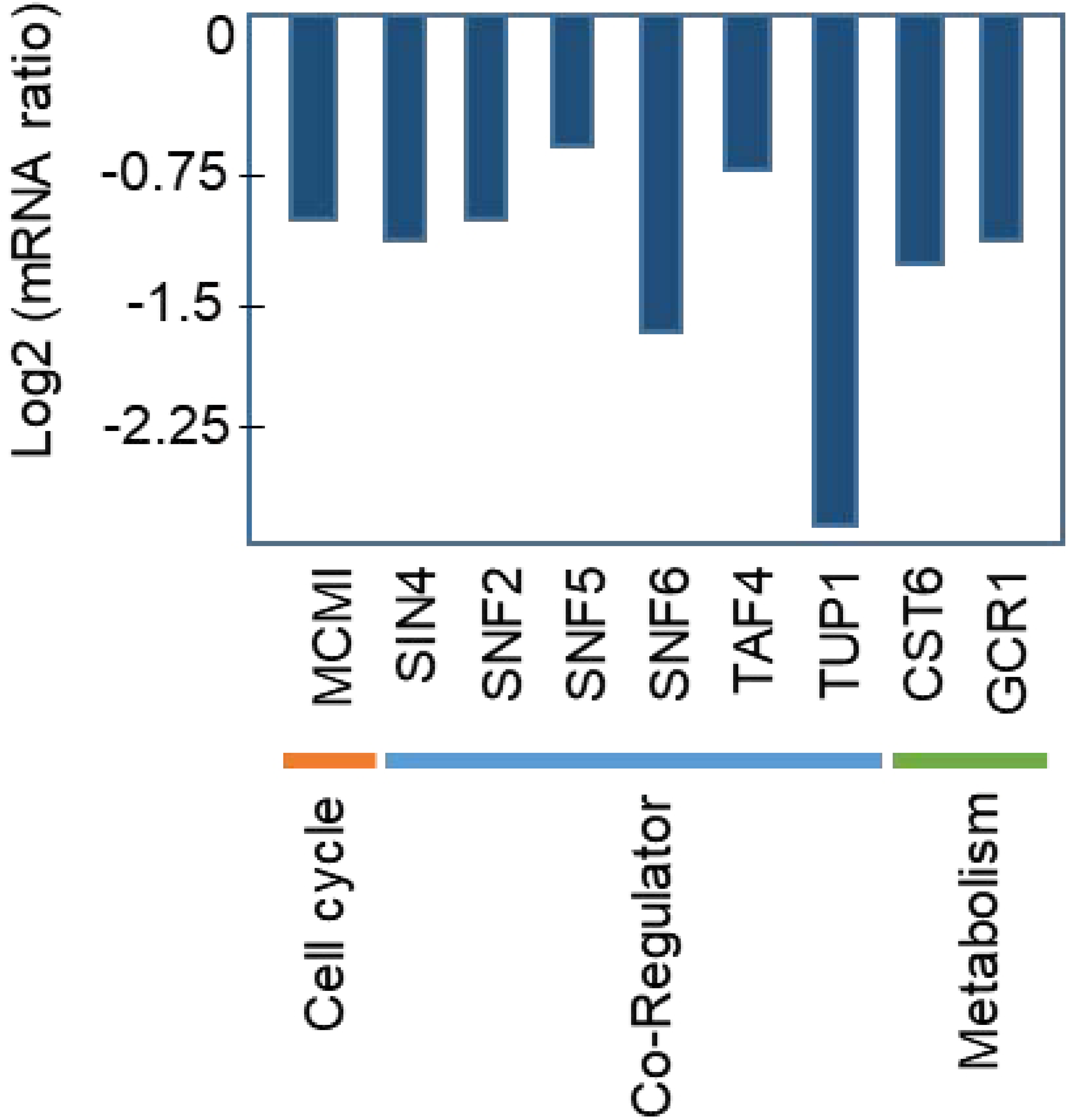
Cluster analysis performed using the Regulator Cluster tool. This analysis groups genes based on the log2 fold change in mRNA expression observed for each gene in each regulator mutant.

**Fig. 3.**
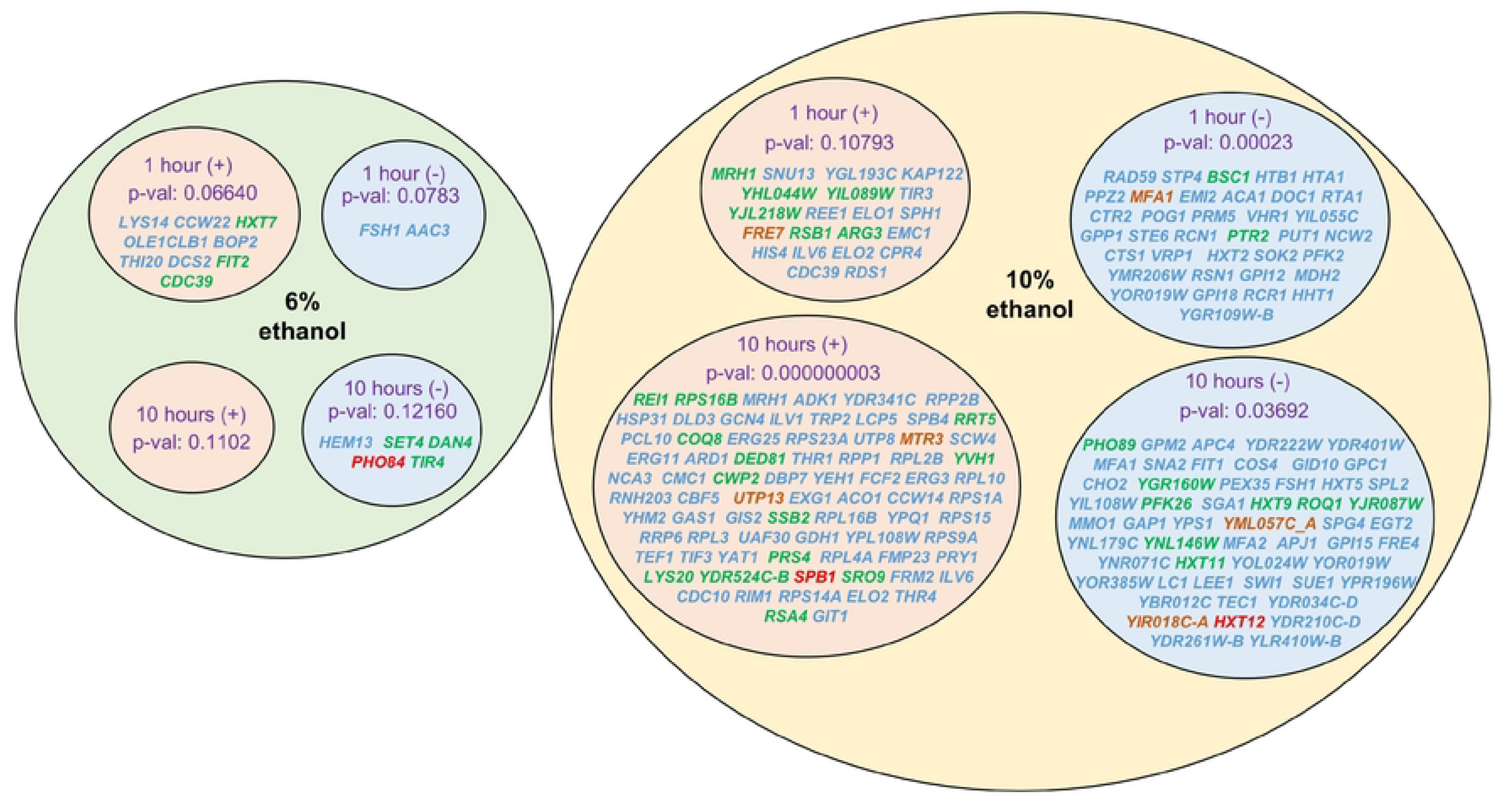
Representation of the differentially expressed genes between strains 2-200-2 and 2-200-2-S4, regulated by *TUP1* in each condition. Genes with a log fold-change >2 are shown in blue, 2–3 in green, 3–4 in orange, and ≥4 in red.

From all the DEGs identified under various ethanol concentration and exposure times (Supplementary Tables 1, 3 and 4), we focused on those regulated by Tup1p with a logFC >1 (Fig.3).Within these DEGs, the genes with the highest logFC (>4), represented in red in Fig. 3, included *PHO84* (a high-affinity inorganic phosphate transporter and low-affinity manganese transporter, downregulated at 6% ethanol and 10 hours exposure), *SPB1* (located on chromosome III, an adoMet-dependent methyltransferase involved in rRNA), and *HXT12* (a sugar transporter). The highest number of DEGs appeared with 10% ethanol presence (Supplementary Table S1 and S5). We conducted gene ontology (GO) analysis on these DEGs, focusing specifically on those regulated by *TUP1*. After 1-hour exposure to 10% ethanol, Tup1p-regulated downregulated genes were primarily involved in chromatin structure. Following 10 hours of exposure to 10% ethanol, upregulated genes were associated with the GO terms cellular component of cytosolic ribosome, and ribonucleoprotein complex. By contrast, downregulated genes were associated with the GO term hexose transmembrane transporter activity.

Notably, we calculated that almost half (45%) of the DEGs were regulated by *TUP1*. Given the diverse functions of these regulated genes, we performed a global pathway enrichment analysis to identify the most significant routes differentiating strains with varying copies of chromosome III.

### Global transcriptional changes

We also examined genes with high differential expression focusing on those enriched in specific pathways according to the KEGG database (Fig. 4). Furthermore, we explored the relationship between these highly expressed pathways and *TUP1*. The most enriched pathway in strain 2-200-2 (three copies of chromosome III) was steroid biosynthesis, which contributes to ethanol resistance by modulating plasma membrane fluidity and is known to be regulated by *TUP1-SSN6*^33^. Also, the tricarboxylic acid (TCA) pathway was highly enriched. Notably, Tup1p plays a key role in regulating metabolic transitions in response to nutrient availability by repressing genes involved in oxidative metabolism, including TCA cycle-related pathways, particularly under glucose-rich conditions^34^. As environmental conditions change, Tup1p repression is alleviated, allowing for the upregulation of metabolic pathways, such as the TCA cycle, to facilitate aerobic respiration and energy production. The next most enriched pathway was porphyrin biosynthesis, which has been linked to *CYC3*, which is highly affected by *TUP1* [35]. In terms of the number of genes affected, the biosynthesis of secondary metabolites was the most enriched pathway, followed by meiosis, cell cycle, biosynthesis of amino acids, and carbon metabolism, containing many genes which are regulated by *TUP1* [26,35].

**Fig. 4.**
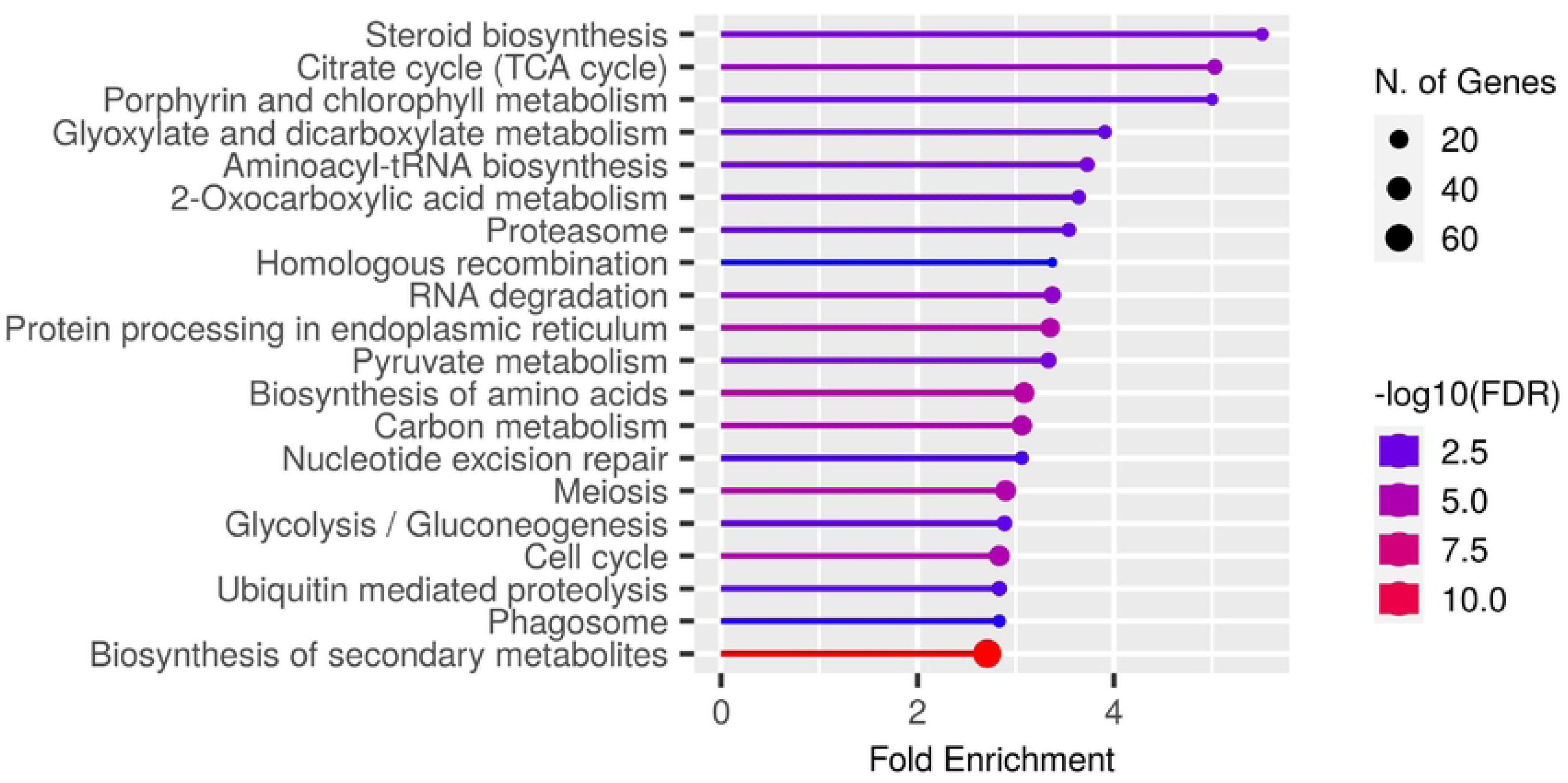
Enriched pathways identified by comparing gene expressions between strains 2-200-2 and 2-200-2-S4 exposed to ethanol. The network visually represents the fold enrichment of each pathway, the number of genes associated with it, and the false discovery rate (FDR). Node size corresponds to the number of genes involved in the respective pathway..

### Deletion of *TUP1* highlights its importance in ethanol response

To investigate the role of *TUP1* copy number in ethanol tolerance, we deleted one copy of *TUP1* (located on chromosome III) from strain 2-200-2 through homologous recombination. Quantitative PCR analysis confirmed that the resulting mutant, 2-200-2Δ*TUP1,* had 2/3 the normal copy number of *TUP1*, indicating successful deletion of one copy. We hypothesized that reducing *TUP1* copy number would decrease ethanol tolerance in 2-200-2Δ*TUP1* mutants, bringing them to a level similar to 2-200-2-S4. To test this, we determined the non-inhibitory concentration (NIC) and minimum inhibitory concentration (MIC) of seven 2-200-2Δ*TUP1* mutants and compared them with 2-200-2 and 2-200-2-S4 strains (Fig. 5). NIC is the concentration above which growth is first visibly affected, while MIC is the lowest concentration of ethanol that completely inhibits visible growth. As expected, 2-200-2Δ*TUP1* mutants exhibited reduced NIC and MIC values compared with strain 2-200-2, aligning with the levels observed with strain 2-200-2-S4. These findings indicate that the increased copy number of *TUP1* on chromosome III contributes to enhanced ethanol tolerance.

**Fig. 5.**
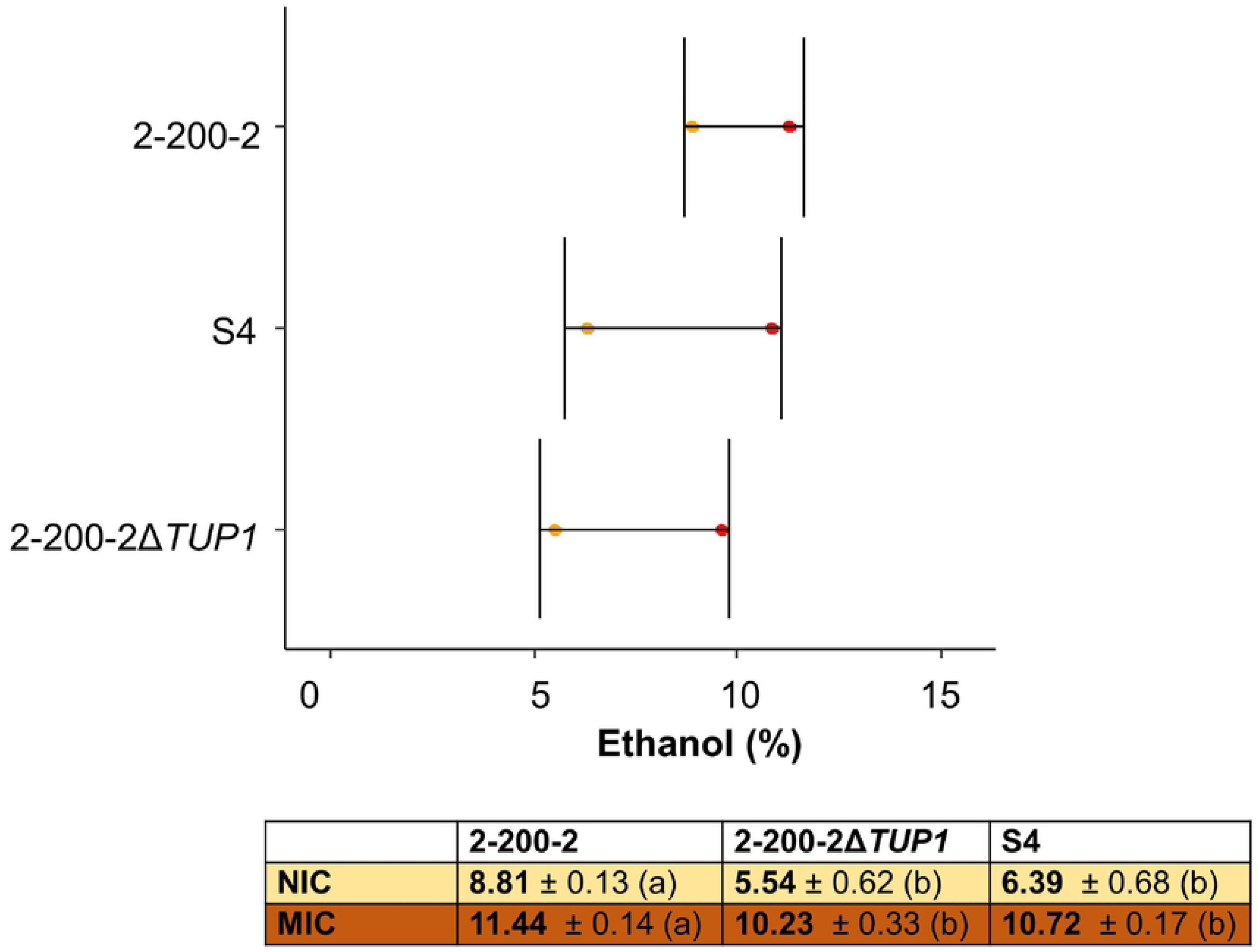
Analysis of NIC and MIC under ethanol exposure in 2-200-2, 2-200-2Δ*TUP1* mutants and 2-200-2-S4. Values were calculated using the area under the curve obtained following the growth curve of the strains under different ethanol concentrations and represent the mean and standard error. Data were analyzed using analysis of variance (ANOVA) followed by Tukey’s HSD post-hoc test, with different groups indicated by the letters "a" or "b".

## Discussion

Ethanol tolerance is a complex trait that remains challenging to fully understand. A common characteristic of ethanol-tolerant yeast strains is the presence of aneuploidy on chromosome III [22]. Previous research has shown that strains with an extra copy of chromosome III exhibit superior tolerance to ethanol than their derivatives with only two copies [21,22].While this suggests a link between chromosome III and ethanol tolerance, the basis for the increased copy number was unclear. In the present study, we compared the expression profiles of two strains with identical genetic backgrounds but differing numbers of chromosome III copies. This allowed us to isolate the effects of the extra chromosome copy. Under normal, ethanol-free conditions, we observed minimal differences in gene expression between the two strains. However, when exposed to high (10%) and low (6%) ethanol concentrations, significant differences in gene expression and activated mechanism emerged. Notably, a substantially larger number of genes were differentially expressed in response to 10% ethanol than to 6% ethanol. These findings align with previous observations that different yeast strains exhibit varying levels of ethanol tolerance, and that these differences may be attributed to diverse underlying mechanisms [35]. Additionally, the time-dependent nature of ethanol resistance [21] suggests that immediate responses to ethanol stress may differ from longer-term adaptations (1 hour *versus* 10 hours in the present study).

A key finding in the present study was that the increased copies of *TUP1* on chromosome III impacted the expression of multiple genes linked to ethanol tolerance. *TUP1,* a conserved transcriptional corepressor, is regulated by glucose depletion and plays a critical role in deacetylation processes, affecting various biological processes including gene expression and transporter activity. It has been shown to influence nucleosome positioning at the *HXT* family [26,30,36] and is essential for quiescent cell survival when acting in concert with *SSN6* [32]. Of note, the Tup1p-Ssn6p complex plays a dual role in *S. cerevisiae*, functioning both as a repressor and an activator depending on cellular conditions [26,28,36]. Under normal conditions, Tup1p primarily functions as a repressor, silencing over 300 genes involved in diverse processes, including carbohydrate metabolism, transport, and stress responses [34]. It is recruited to specific promoters by DNA-binding proteins such as Sko1p, repressing transcription through both chromatin-dependent and chromatin-independent mechanisms [29,30]. Interestingly, the repression of genes encoding chromatin structural components, observed after short-term ethanol stress (1 hour, 10% ethanol), may facilitate broader transcriptional changes. During stress conditions, including ethanol or osmotic stress, the function of Tup1p shifts [35,37], as the cell undergoes a complex reorganization of gene expression, characterized by increased activation of genes involved in the cytosolic ribosomal complex. Tup1p is implicated in recruiting coactivators such as *SAGA*, *SWI/SNF*, and mediators to promoters, suggesting a switch from repressor to activator [27]. The increased activation of cytosolic ribosomal genes, observed after prolonged stress (10 hours), likely reflects a heightened need for protein synthesis to produce stress-mitigating factors. Concurrently, the cell exhibits differential regulation of sugar transporters, finely tuned to ethanol stress levels. At higher concentrations (10% ethanol), our results showed that sugar transporter genes are repressed, while at lower levels (6% ethanol), the high-affinity transporter *HXT7* is activated. This strategy optimizes resource utilization [38-40]. Tup1p is likely involved in this dynamic regulation, given its role in repressing carbohydrate metabolism and transport genes under normal conditions [34]. During osmotic stress, activation of the *HOG* pathway leads to Hog1p phosphorylation, causing the disassembly of the Tup1p-Ssn6p-Sko1p repressor complex and converting Sko1p from a repressor to an activator^27^. During hyperosmotic stress, Tup1p undergoes rapid and transient sumoylation at lysine 270 (K270), which is crucial for its functional switch [37,41]. This sumoylation, dependent on Cyc8p association, is a major contributor to the overall SUMO conjugation observed during stress^13,41^. The duration of Tup1p sumoylation is regulated by the Hog1p MAPK pathway, with prolonged sumoylation occurring in *HOG1*Δ cells [37]. This sumoylation, in combination with other post-translational modifications and protein interactions, enables Tup1p to activate stress-response genes at specific promoters, such as *GPD1* [37]. As the cell adapts and accumulates glycerol, Tup1p-Cyc8p is desumoylated, resetting the stress response. This dynamic regulation allows *S. cerevisiae* to rapidly adapt to environmental changes [37,42]. Aneuploidy, specifically chromosome III duplication, can further influence this stress response. Increased copy number of *TUP1* can alter gene expression patterns through gene dosage effects impacting ethanol tolerance. Changes in copy number can lead to varying levels of gene expression, potentially leading to different levels of transcription [43]. Additionally, chromosomal duplication serves as an initial evolutionary defence mechanism, enabling organisms to withstand sudden and intensive selective pressures [24].

In conclusion, our study reveals that *S. cerevisiae* employs a sophisticated adaptation mechanism to tolerate ethanol stress, centered around the transcription factor *TUP1*. We found that the copy number of *TUP1*, located on chromosome III, significantly influences ethanol tolerance, as evidenced by the decreased ethanol tolerance in 2-200-2Δ*TUP1* mutants. Ethanol-tolerant strains exhibit increased copy number of chromosome III, likely amplifying *TUP1* expression. This chromosomal duplication serves as an initial evolutionary response to sudden, intense selective pressures. The ability of *TUP1* to rapidly switch between activator and repressor roles through sumoylation allows for rapid adaptation to changing environments. While *TUP1* is a general stress regulator, our study highlights its specific importance in ethanol tolerance. We identified numerous genes differentially regulated by *TUP1* at high ethanol concentrations in ethanol-tolerant strains, emphasizing its crucial role in orchestrating the cellular response to ethanol stress. This comprehensive evidence underscores the complex interplay between chromosomal duplication, gene regulation, and adaptive responses mediated by *TUP1* in ethanol tolerance, providing insights into the evolution of stress resistance in yeast.

## MATERIALS AND METHODS

### Strains and media

*S. cerevisiae* strain 2-200-2 is a haploid-derived diploid from strain FY5 possessing three copies of chromosome III [21]. Strain 2-200-2-S4 is a derivative of 2-200-2 with one copy of chromosome III deleted [22]. Yeast strains on a 2-200-2 background but with *TUP1* deleted were created in the present study, and explained later. Yeast cells were cultured in Glucose Peptone Yeast Extract (GPY) medium, which contained 20 g/L of glucose, 20 g/L of bacteriological peptone, and 10 g/L of yeast extract. For solid media, 20 g/L of agar was added.

### Fermentation for sample extraction

Strains 2-200-2 and 2-200-2-S4 were inoculated in GPY medium and grown overnight. The cell suspensions were then adjusted to an initial OD_600_ of 0.2 in 1L Erlenmeyer flasks containing GPY medium with 0%, 6%, or 10% ethanol. This setup was performed in triplicates for each strain and condition. The flasks were incubated at 28°C with orbital agitation at 150 rpm. At 1 and 10 hours post-inoculation, approximately 10^8^ cells were collected from each flask. These samples were immediately frozen in liquid nitrogen and subsequently stored at -80°C for further analysis.

### RNA sequencing

RNA was isolated using the High Pure RNA Isolation kit (Roche Applied Science, Mannheim, Germany) and oligo (dT) mRNA purification. RNAseq libraries were generated using the TruSeq Stranded mRNA Library Preparation Kit (Illumina Inc., San Diego, CA). The information about the sequences area available in BioProject ID PRJNA1153811.

### RNASeq data analysis

Reads were mapped to the *S. cerevisiae* reference strain S288C using Bowtie2 (Bowtie2 v.2.2.9, local alignment mode) [44]. Alignments were then compressed and sorted using SAMtools v.1.4.1 [45].Finally, read counts for each gene were determined using HTSeq-Count (HTSeq-0.6.1p1, parameters −m union –a 10) [46].

PCA was performed on the gene expression data using the “dds” function from the DESeq2 R package, which normalizes and adjusts the sample counts to a negative binomial distribution, and then applies a variance stabilizing transformation function. The DEGs between pairs of conditions were identified using the contrast function within DESeq2. Fold changes and corresponding p-values were determined for each DEG.

The intersection of gene sets was visualized using Venny [47]. GO enrichment analysis was performed using PANTHER19.0 [48] with Fisher’s exact test and a false discovery rate (FDR) threshold of P < 0.05 (Benjamini-Hochberg correction). Pathway enrichment analysis was performed using ShinyGO [49] with the list of DEGs. Finally, potential transcription factors responsible for the observed differential gene expression were identified using the YEASTRACT+ [32] and DB [33] databases

### Mutant construction

To disrupt the *TUP1* gene in strain 2-200-2, we employed the LiAc/SS Carrier DNA/PEG transformation method [50]. The KanMX cassette was amplified from plasmid pUG6 using primers incorporating homology regions flanking the *TUP1* open reading frame. Transformants were selected on GPY medium supplemented with G418. After three days, colony PCR was performed with genomic DNA extracted using the LiAc method [51] and specific primers (Supplementary Table S6) to confirm the insertion of the cassette. Since 2-200-2 harbors three copies of *TUP1*, the disruption could potentially result in the deletion of one or two copies of *TUP1*. We used qPCR analysis to determine the remaining copies of this gene.

The primers were designed to amplify an internal sequence of *TUP1* and also *ABP1*, which is also found in chromosome III near *TUP1*. *ABP1* has previously been used to confirm chromosome copy number in *Saccharomyces* [52].

DNA from all strains was extracted in quintuplicate using a modification of a published extraction method [53]. The extracted genomic DNA was then treated with RNAse A and phenol-chloroform to remove RNA and proteins. DNA concentration was quantified by fluorimetry using a Qubit™ 4 Fluorometer with the Qubit™ 1× dsDNA BR Assay (Invitrogen, Carslbad, CA), and samples were diluted to 10 ng/µl. Real-time qPCR was performed with 2 µl of each sample, primers specific for *TUP1* and *ABP1* (Additional information Table S1), and the NZYSpeedy qPCR Green Master Mix (2×) (NZYtech, Lisbon, Portugal) on a LightCycler® 480 System (Roche Applied Science). Three technical replicates were conducted for each of the five DNA extractions from every strain. The results from the qPCRs were statistically analyzed to calculate the average expression levels of the target gene *TUP1* and the reference gene *ABP1*. The ratio *TUP1/ABP1* was then compared between the wild strains (FY5, 2-200-2, and 2-200-2-S4) and their respective mutants to assess whether the copy number decreased. Finally, the mutant stains were sequenced to confirm the introduced changes.

### Ethanol tolerance

*Saccharomyces* strains were cultured overnight in 1 ml of GPY. Subsequently, cultures were centrifuged to separate the cells, washed, and resuspended in PBS for 2 hours. Cell density was determined using a DeNOVIX CellDrop FL (Wilmington, DE) cell counter. Based on these measurements, the appropriate volume of each strain was calculated to achieve a final concentration of 4×10^7^ cells/ml. From this suspension, 11 µl of each strain was inoculated (in triplicate) into 96-well plates. Each well contained 220 µl of minimal YNB medium and different ethanol concentrations. To prevent evaporation, 100 µl of Vaseline oil was overlaid on the culture medium.

Yeast growth was monitored by measuring the OD_600_ using a Stacker Microplate Handling System connected to a SPECTROstar Omega plate reader (BMG LABTECH, Ortenberg, Germany). The microplates were incubated within a 25°C, 70% humidity chamber equipped with an orbital shaker. Microplates were shaken for 20 seconds at 400 rpm every hour. Growth curves were analyzed using the Growth Curve Analysis Tool (GCAT) [54] and the best-fitting model (Richards, Gompertz, or logistic sigmoid). Subsequently, growth parameters (lag time, growth rate, and asymptotic growth value) were determined using reparametrized formulas described by Zwietering et al. (1990) [55]. To quantify overall growth, the area under the curve (AUC) was obtained by integrating the curve from 0 to 70 hours (final time). The relative growth of each yeast strain at different ethanol concentrations was determined by calculating the fractional area (fa). This is calculated by:

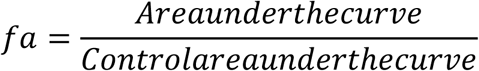

A modified Gompertz function was used to relate the fractional area (y) to the log of ethanol concentration (x) [56]:

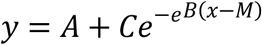

In this formula, A is the lower asymptote of y, B is the slope parameter, C is the distance between the upper and lower asymptote and M is the log concentration of the inflection point. The values for NIC and MIC are described as the intersection of the lines y = A+C and y = A, with the equation of the line tangential to the point (M) respectively.

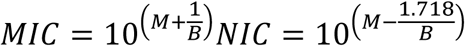

The values of A, C, B, and M can be calculated using a non-linear fitting procedure, and NIC and MIC are determined.

## DECLARATIONS

### Availability of data and materials

The information about the sequences area available in BioProject ID PRJNA1153811.

